# Dimensions of neighborhood tracts and their associations with mental health problems

**DOI:** 10.1101/518258

**Authors:** Katherine L. Forthman, Hung-wen Yeh, Rayus Kuplicki, Martin P. Paulus

## Abstract

**Objective:** Neighborhood characteristics can have profound effects on resident health. The aim of this study was to use an unsupervised learning approach to reduce the multi-dimensional assessment of a neighborhood using American Community Survey (ACS) data to simplify the assessment of neighborhood influence on health.

**Method:** Multivariate quantitative characterization of the neighborhood was derived by performing a factor analysis on the 2011-2015 ACS data. The utility of the latent variables was examined by determining the association of these factors with poor mental health measures from the 500 Cities Project 2017 release.

**Results:** A five-factor model provided the best fit for the data and the latent factors quantified the following characteristics of the census tract: (1) affluence, (2) proportion of singletons in neighborhood, (3) proportion of African-Americans in neighborhood, (4) proportion of seniors in neighborhood, and (5) proportion of noncitizens in neighborhood. African-Americans (*R*^*2*^ = 0.67) in neighborhood and Affluence (*R*^*2*^ = 0.83) were strongly associated with poor mental health.

**Conclusions:** These findings indicate the importance of this factor model in future research focused on the relationship between neighborhood characteristics and resident health.

## Introduction

There is strong evidence that certain characteristics of the neighborhood are associated with both the physical and the mental health of its residents [1–7]. However, the strength of this effect and the qualities of the neighborhood environment that are related to the effect are unclear. Contradictory results in past studies relating neighborhood characteristics and health indicate that the environmental effect remains elusive [8]. Thus, objectively delineating the dimensions of a neighborhood and evaluating their relationship to mental health could have important implications for future research linking social factors to the biological processes underlying psychiatric disorders.

Past research has not yet fully elucidated the relationship between mental health and neighborhood environment for several reasons. First, most studies only evaluate the effects of one neighborhood characteristic—commonly neighborhood socioeconomic status (NSES) or racial composition—rather than looking at a broader range of variables describing the neighborhood. There is an emerging consensus that NSES is correlated with physical and mental health, however the lack of more information about the nature of this relationship leaves no hint of how NSES might be linked to these health effects. Second, amidst the current literature are many different approaches in ascertaining a population, extracting neighborhood measures, focusing on specific indices, or selecting a subset of the population. A comprehensive approach has been lacking and could provide a crucial step forward to identify specific environmental factors influencing rates of poor mental health.

The American Community Survey (ACS) a household survey conducted by the U.S. Census. It covers a vast number of statistics and includes a substantial sample of the U.S. population. Therefore, it is an ideal dataset for studying demographics. In order to use as many of the statistics provided by the ACS as possible, we used a latent variable approach to arrive at a multivariate quantitative characterization of the neighborhood. This method gave us the opportunity to objectively select variables to include in our analysis based on their contribution to the variance between neighborhoods. We assumed that these statistics are linked by underlying, latent variables. This allowed us to work with a smaller number of variables without sacrificing any data.

Few studies have used such a method for the characterization of neighborhoods. Miles et al. describes a method for measuring NSES using factor analysis and the ACS that can be explored longitudinally [9]. Their method aims to find significant neighborhood characteristics based on factor invariance. Another study by Li et al. combines factor analysis and cluster analysis for a multivariate-structural approach to characterizing neighborhoods [10]. These studies were used as a template for the method described in this paper. This study aimed to use a latent variable approach to identify factors that comprehensively quantify the neighborhood characteristics. We used the 2011-2015 ACS data and applied a factor analysis method after appropriately transforming selected variables. Moreover, we examined the influence of these latent variables on mental health by comparing the factors to data from the 500 Cities project 2017 release [11]. The 500-cities dataset includes 27,199 tracts. We performed this analysis in order to explore if one or more factors would be significantly associated with proportion of individuals in a neighborhood with at least 14 “bad mental health days” as ascertained by the Centers for Disease Control and Prevention (CDC).

## Methods

### Sample

Neighborhood data were obtained from the ACS. The ACS is a national survey that uses continuous measurement methods based on a series of monthly samples to produce annual estimates for the same small areas (census tracts and block groups). The ACS was inaugurated in 2005. Each year, ~3.54 million addresses are surveyed across the country. This sample is sufficient to provide reliable 1-year estimates for geographical areas with a population greater than 65,000; for areas with smaller populations, the sample needs be accumulated over several years to achieve reliable estimates (3-year and 5-year estimates for areas with population larger than and no more than 20,000, respectively) [12]. In this study, neighborhood data were obtained at tract-level from the 5-year estimate spanning 2011-2015 [13].

This study aimed to use data at the smallest geographic division possible for a fine-grained view of living environment. The two smallest geographic divisions available in the ACS dataset are block-groups and tracts. On average, a block group is 1/3 the size of a tract. The factor analysis was performed on both levels and the resulting factors were largely the same (Comparison to Block Group, Supplementary Methods). Though larger, the tract level factors were chosen because there were variables of interest that were not available at the block group level (Fig S1). These variables included disability status, citizenship status, and mobility. Additionally, the tract-level factors have a smaller margin of error.

There are 72,424 census tracts in the United States that are within city and state boundaries. Tracts consist of areas with a population between 1,200 and 8,000 individuals, are primarily defined by population density, and are delineated by visibly identifiable features, such as highways, roads, or rivers. Both home addresses and group quarters were sampled from the tracts. Group quarters are places where a group of people live together in a place which is owned or managed by an entity that provides housing or services to residents; for example, nursing homes, college dormitories, and homeless shelters are all group quarters. Approximately 2.5% of the expected population inhabiting group quarters was sampled [12].

### Data extraction and variable selection

ACS variables are organized into tables, which are organized by content and format [14]. Not all tables were used in this analysis. A description of the excluded tables and the reason for exclusion is provided in the supplementary material (Table S1). 37 tables were included in the analyses and contained 461 measures describing tract characteristics of age, race/ethnicity, citizenship, nativity, mobility, means of transportation to work, household type, marriage status, education level, disability status, income, employment status, home type, housing cost, and residential tenure. Among these 461 measures, 215 were removed due to redundancy or low variability. Redundancy was defined as variables that were sufficiently represented by another variable (for example, the female population is redundant because it is the inverse of the male population). Low variablility was defined as a low coefficient of variation across tracts. A flow chart explaining how the dataset was reduced is given in Fig S2. The remaining 246 statistics describing strata or subgroups (e.g. age groups, gender, education levels) of a tract were combined to form single statistics (Feature Selection, Supplementary Methods). This selection process led to a final number of 39 measures for subsequent analyses.

These measures were subjected to a heuristic, data-driven transformation approach to approximate Gaussian distributions as close as possible (Transformation, Supplementary Methods). Missing values were imputed by the weighted average of 10-nearest neighbors after transformation.

### Factor analysis

A total of 39 transformed and/or imputed measures were then entered into an exploratory factor analysis to investigate the underlying latent variable structure. Factor loadings were estimated by the minimum residual method, and oblimin rotation was applied to improve interpretation. We explored a range between 1 and 12 factors and chose the factor number based on Kaiser’s rule (i.e. keeping factors with eigenvalues at least 1), a scree plot, the amount of total variance explained from each model produced, and the interpretability of the factor structure. The chosen factor model is depicted in Fig 1. The factor structures of the 4- and 6-factor models are shown in Figs S3 and S4 for comparison purposes. The stability of the final factor model was examined by 2,000 bootstrapped samples and the standard error was calculated for each loading of each variable within each factor. Finally, the factor scores were computed for all U.S. tracts.

**Fig 1.**
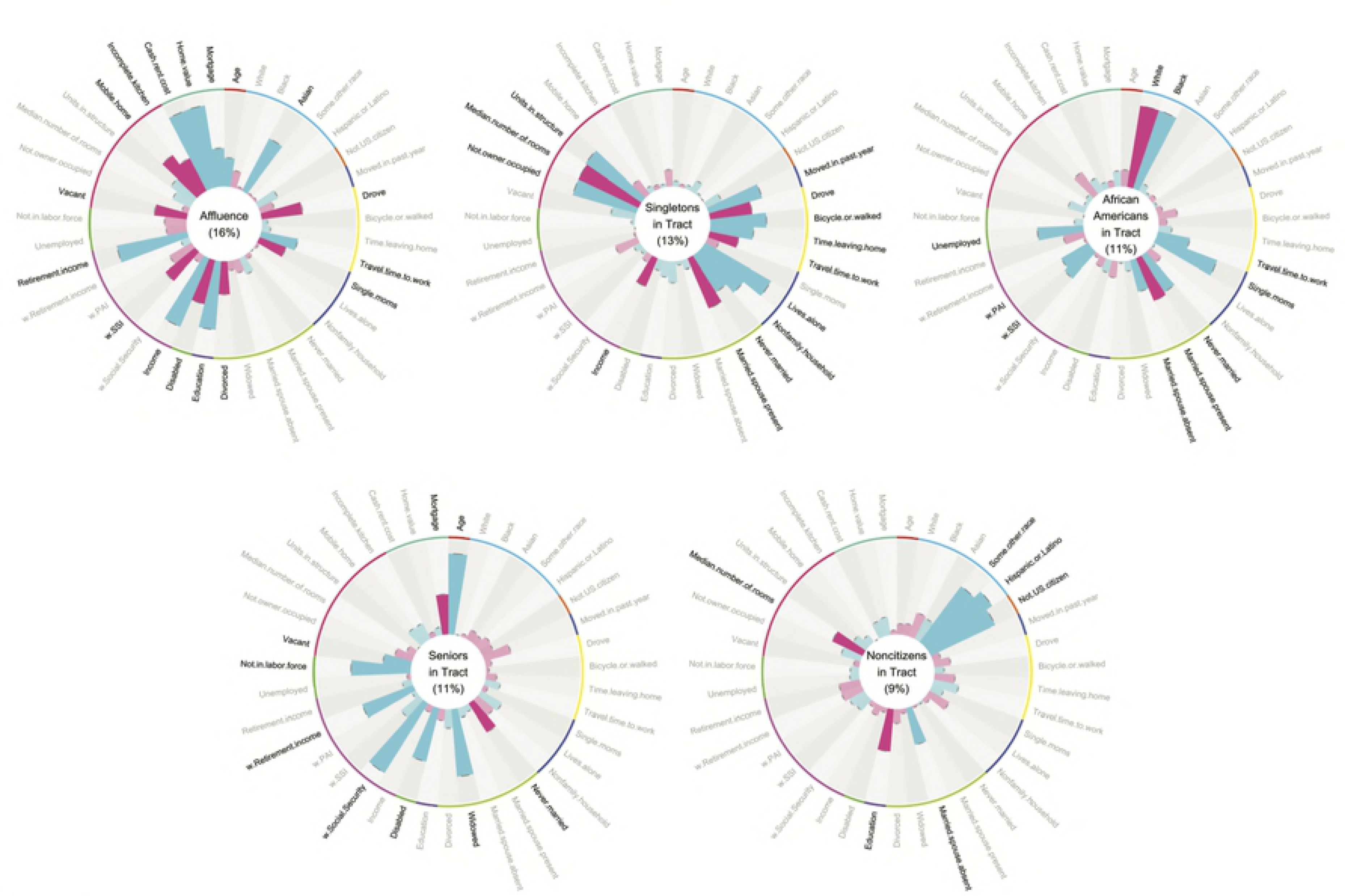
The Factor Structure. Each circular barplot is a visual representation of a single latent factor. The name of the factor is in the center of the plot. Each bar represents the loading of an input variable to the factor. Blue bars indicate a positive loading, while pink bars indicate a negative loading. Variables with loadings > 0.3 to the factor are highlighted. Input variables are grouped by type with the colored lines around the edge of each plot. These groups (starting from the top, moving clockwise) encompass age, race and ethnicity, nativity and citizenship, mobility, transportation to work, household type, marital status, education level, disability status, income, employment, residential conditions, and tenure. A larger version of these plots is given in the supplement (S2).

### Relationship between neighborhood latent variables and mental health

The factor scores for each census tract were merged with data from the 500 Cities Project, which provides tract-level mental health data for 27,204 tracts [11]. This project used Small Area Estimation (SAE) to estimate prevalence of health issues. The SAE was performed on datasets managed by the CDC, including the Behavioral Risk Factor Surveillance System (BRFSS) [15]. The BRFSS was conducted by a telephone survey interviewing approximately 400,000 adults across the United States and its territories [16]. Iterative proportional fitting was used to weight statistics by age, gender, race and ethnicity, and geographical region [17]. For the purposes of this study, we focused on only one question in the BRFSS: “*Now thinking about your mental health, which includes stress, depression, and problems with emotions, for how many days during the past 30 days was your mental health not good?*” The tract-level SAE from this question provided an estimate of the proportion of individuals ≥18 years old within a tract who responded that they had ≥14 bad mental health days. These estimates were linked to the neighborhood factor scores and their associations were investigated descriptively by smoothing splines.

### Software

The statistical software R [18] was used for all data extraction, analyses, and the generation of all figures. The R code for this manuscript is available as a supplement. ACS data were obtained through the R package *acs* [19]. The *e1071* (31) and *scales* (32) packages were used for transformation, the *DMwR* package for imputation, and the *psych* package [20] for factor analysis.

## Results

### Exploratory factor analysis

The scree plot shows that a factor model with up to eight factors had eigenvalues greater than 1.0 and an ‘elbow’ at 5 factors (Fig S5). Together, these factors accounted for 60% of the variance and reproduced 0.98 of the off-diagonal elements of the sample correlation matrix, and 0.04 root-mean square of residuals (RMS). Fit statistics of the 12 factor models explored are given in Table S2. The five factors were labeled based on the variables with the strongest absolute loadings as: (1) Affluence, (2) Singletons in Tract, (3) African-Americans in Tract, (4) Seniors in Tract, and (5) Noncitizens in Tract (Fig S6). Affluence, which accounted for 16% of the variance, showed greatest loadings from tract statistics relating to NSES, such as income (0.79 for Income in the circle plot) and education (0.73 for Education). Singletons in Tract, which accounted for 13% of the variance, demonstrated strong loadings from the proportion of people living alone (0.81 for Lives.alone), the average number of housing units per structure (0.72 for Units.in.structure), and the proportion of homes in a tract not occupied by their owner (0.70 for Not.owner.occupied). African-Americans in Tract, which accounted for 11% of the variance, was positively correlated with the proportion of black population (0.87 for Black) and inversely correlated with the proportion of white population (−0.87 for White). This factor was also highly correlated to proportion of single moms (0.69 for Single.moms), a lack of married couple family homes (−0.49 for Married.spouse.present, 0.46 for Never.married), the unemployed population (0.49 for Unemployed), and the proportion of people living on government assistance (0.37 for w.SSI, 0.34 for w.PAI). Seniors in Tract, which accounted for 11% of the variance, was primarily related to age (0.85 for Age) and the proportion of the population receiving Social Security Income (0.87 for w.Social.Security). Noncitizens in Tract, which accounted for 9% of the variance, was strongly related to the proportion of certain racial and ethnic minorities (0.74 for Some.other.race, 0.83 for Hispanic.or.Latino) as well as the population of U.S. citizens (0.76 for Not.US.citizen).

The oblique rotation procedure left the factors correlated (Fig S7): African-Americans in Tract was correlated with Noncitizens in Tract (r = 0.36), Singletons in Tract (r = 0.33), and Affluence (r = −0.29); Seniors in Tract was correlated with Noncitizens in Tract (r = −0.26) and Affluence (r = −0.21). Least correlated were Noncitizens in Tract and Affluence (r = −0.08).

### Associations between neighborhood factors and prevalence of poor mental health

The prevalence of individuals in a tract with 14 or more days of bad mental health appeared to be most related to Affluence, followed by African-Americans in Tract, Noncitizens in Tract, Singletons in Tract, and least related to Seniors in Tract (Fig 2). There was an obvious inverse relationship between the bad mental health measure and affluence of tracts for all states (median R-square 0.83 and inter-quartile range (IQR) between 0.80 and 0.86, Fig S8). There also existed monotone, increasing trends between the health measure and the two factors African-Americans in Tract (median and IQR *R*^*2*^: 0.67 (0.58, 0.74)) and Noncitizens in Tract (median and IQR *R*^*2*^: 0.49 (0.35, 0.67)), despite higher variability in trends across states for the latter. Concave trends appeared between the bad mental health outcome and Singletons in Tract for most states (median and IQR *R*^*2*^: 0.17 (0.11, 0.24)). The uniform relationship for tracts with a lower Singletons in Tract score indicates that neighborhoods with fewer singletons tend to have lower rates of poor mental health. As the factor score increases, however, rates of mental health become more variable, indicating there is no relationship between mental health and a higher Singletons in tract score. Seniors in Tract showed different patterns across states, with a mixture of positive and negative, linear and concave trends (median and IQR *R*^*2*^: 0.05 (0.02, 0.11)).

**Fig 2.**
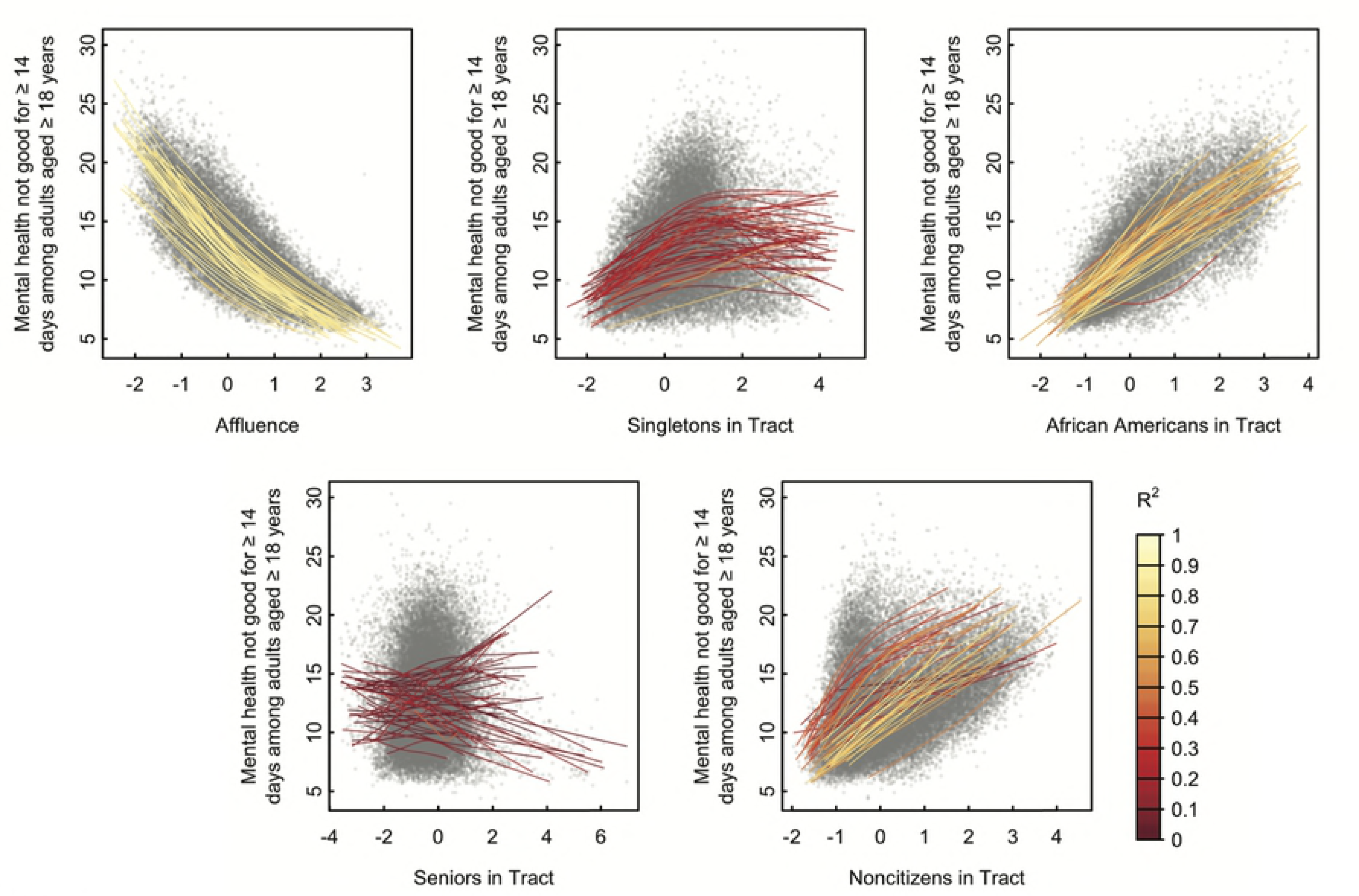
Relationship Between Factors and Mental Health. Proportion of residents over 18 who have experienced ≥ 14 days of bad mental health during the past 30 days from the 500 Cities Project vs. neighborhood factor scores. Each point on the plot represents a single tract. A separate cubic spline (colored curve) was fit to tracts of each state.

## Discussion

This study aimed to quantify neighborhood characteristics using a latent variable approach performed on census data and to determine the utility of these latent variables by relating them to mental health outcomes. There were two main results. First, five factors, Affluence, Singletons in Tract, African-Americans in Tract, Seniors in Tract, and Noncitizens in Tract, accounted for 60% of the neighborhood tract variance and provided a multidimensional assessment of census tracts. Second, two of the five factors were shown to be strongly related to tract-level descriptors of poor mental health: Affluence, and African-Americans in Tract. Taken together, this study shows that census tracts can be robustly quantified using five dimensions and that some of these latent variables are strongly associated with tract-level mental health status.

Several studies have described the relationship between neighborhood characteristics and mental health [8, 21, 22]. However, to our knowledge, no previous study has used a factor analysis approach to extract a multi-dimensional set of latent variables for the characterization of neighborhoods. A recent study by Hu et al. was published showing the relationship between NSES and health [23]. However, the results of these analyses are limited because–as in many other papers–only NSES was examined as a predictor of health while the possible influence of other neighborhood characteristics was ignored. Additionally, the Area Deprivation Index (ADI) used in this paper, developed by Singh [24], was developed based on a single-factor analysis using 17 socioeconomic indicators selected by Singh from the 1990 U.S. census, however time-invariance of the results of the factor analysis was not tested. It is important that this statistic was not tested for time-invariance, as assuming the factor structure is consistent over time may lead to biased results if time invariance of the factor structure does not hold [25]. An index should be based on data in the relevant time period if time-invariance has not been demonstrated. A factor model can be easily calculated for any 5-year period after 2005 using the methods we describe in this paper.

Our results indicate that our factors representing Affluence and African-Americans in Tract are most predictive of mental health rates in a neighborhood. Furthermore, results indicate the pattern each relationship follows. The relationship between African-Americans in Tract and mental health appears linear, while the relationship between Affluence and mental health appears to follow an exponential decay. Our four most explanatory factors (Affluence, African-Americans in Tract, Singletons in Tract, and Noncitizens in Tract), have all been explored to some extent in past research. Affluence appears to be synonymous with such measures as socioeconomic status, economic disadvantage, and neighborhood deprivation as described in several previous studies [8]. African-Americans in Tract and Noncitizens in Tract have also been explored in a few past studies as ‘racial congruence’ or ‘ethnic diversity’, etc. [8]. Even Singletons in Tract is representative, to an extent, of residential mobility or neighborhood stability [8]. The only factor not explored in previous studies was the elderly population, which we have shown to be uncorrelated with rates of mental health. The relationship of our factors to interests of previous studies indicate that our factors are intuitively as well as objectively descriptive of neighborhoods.

## Limitations

The variables used in this study were limited to those collected by the U.S. census. Consequently, there are some neighborhood characteristics shown in previous studies to be related to mental health that are not included in this study. For example, this study does not include walkability, neighborhood disorder, social factors, neighborhood hazards, the built environment, or the service environment [8]. Even with these limitations, the ACS dataset serves as a reliable source for neighborhood statistics. These statistics include responses from millions of households across the U.S., the data are collected consistently over time, and the statistics cover a broad range of characteristics. Additionally, neighborhood characteristics based on subjective resident response may be biased and misleading. For example, perception of neighborhood conditions has been shown to be significantly correlated to rates of depression [8], but there is no assurance that this relationship does not simply depict the poor outlook of those with depression. Additionally, there exist some challenges from an analysis standpoint. The indeterminacy problem is a well-known issue with factor analysis [26, 27]. Factor analysis results in factors that must be subjectively defined. This has always been a fundamental problem of factor analysis. However, the circle plots clearly depict what the factors represent. Additionally, this problem is superseded by the utility of the latent variables over the raw data. The latent variable structure, though vague, simplifies interpretability drastically.

## Future directions

The influence of these factors at an individual level is still unknown. Jones et al. make a point that people experience substantial segregation across a range of spaces, such as areas of work or recreation, in their daily lives [28]. The extent to which an individual’s neighborhood characteristics affect their mental health must be explored in future longitudinal studies. The 500-Cities data contains data on physical health as well as mental health. An exploration on the relationship between the factors and the other statistics in the 500-Cities dataset is given in Fig S9. It is clear from this plot that the factors relate to more than just mental health.

## Conclusion

Neighborhood factors based on census data provide comprehensive, objectively derived neighborhood characteristics. To our knowledge, our work takes into consideration a variety of neighborhood statistics not previously explored, while remaining simple and highly interpretable. We intend that these factors may be used to further explore the relationship between living environment and mental health. Our findings show that neighborhood characteristics are strongly related to mental health, indicating the importance of the factor model in future research focused on the influence of neighborhood characteristics on mental health.

---

**Fig S1.**
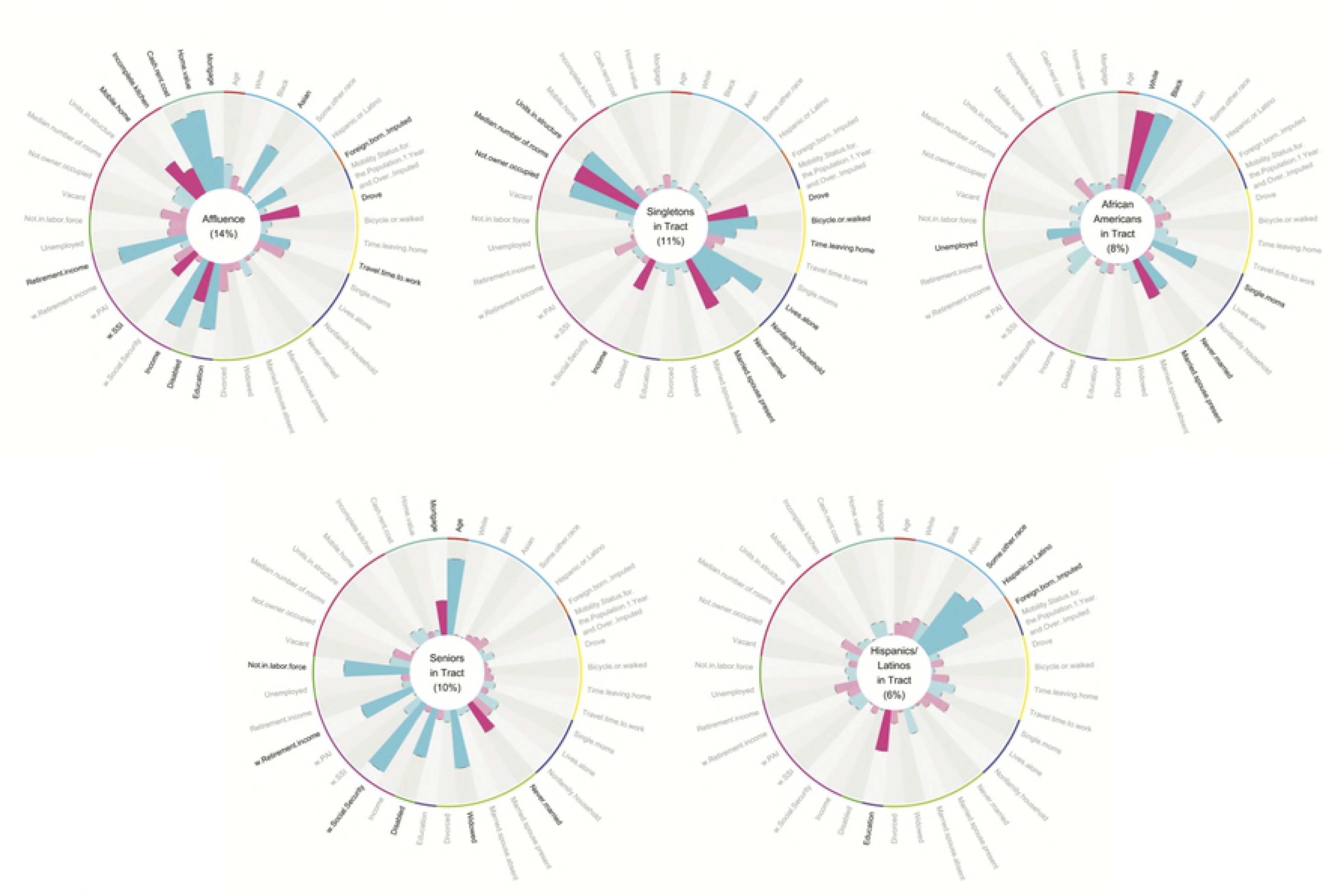

**Fig S2.**
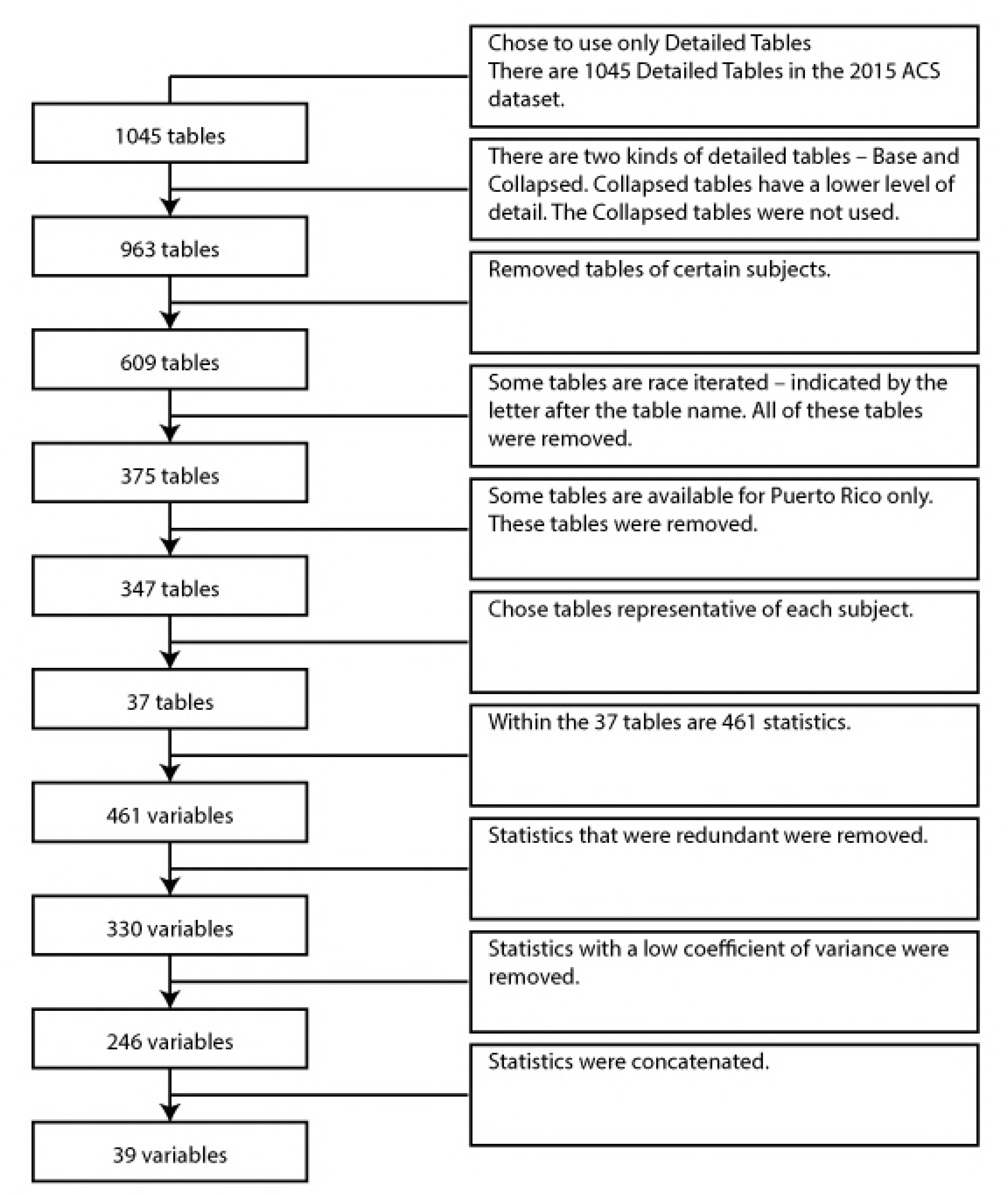

**Fig S3.**
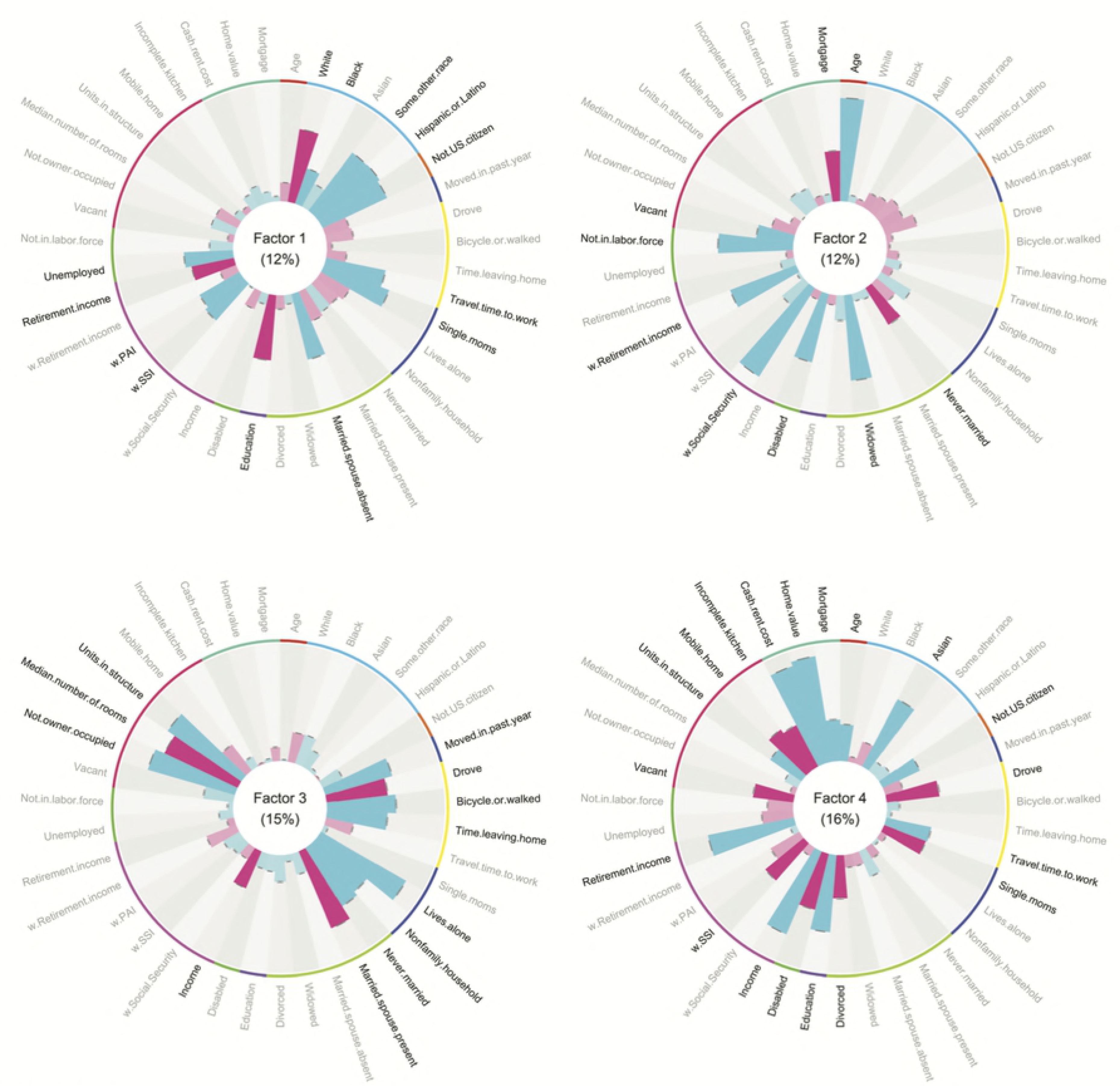

**Fig S4.**
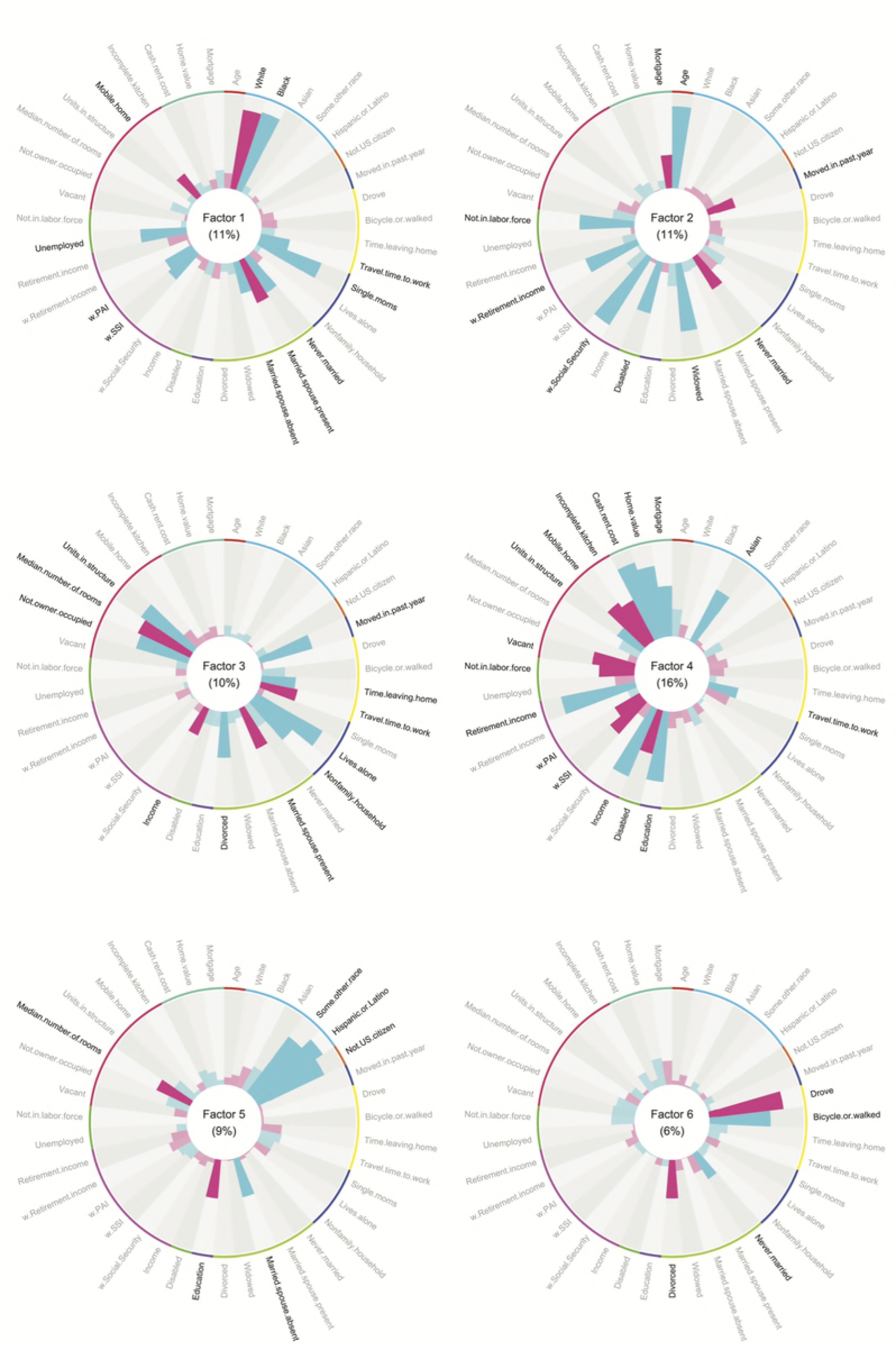

**Fig S5.**
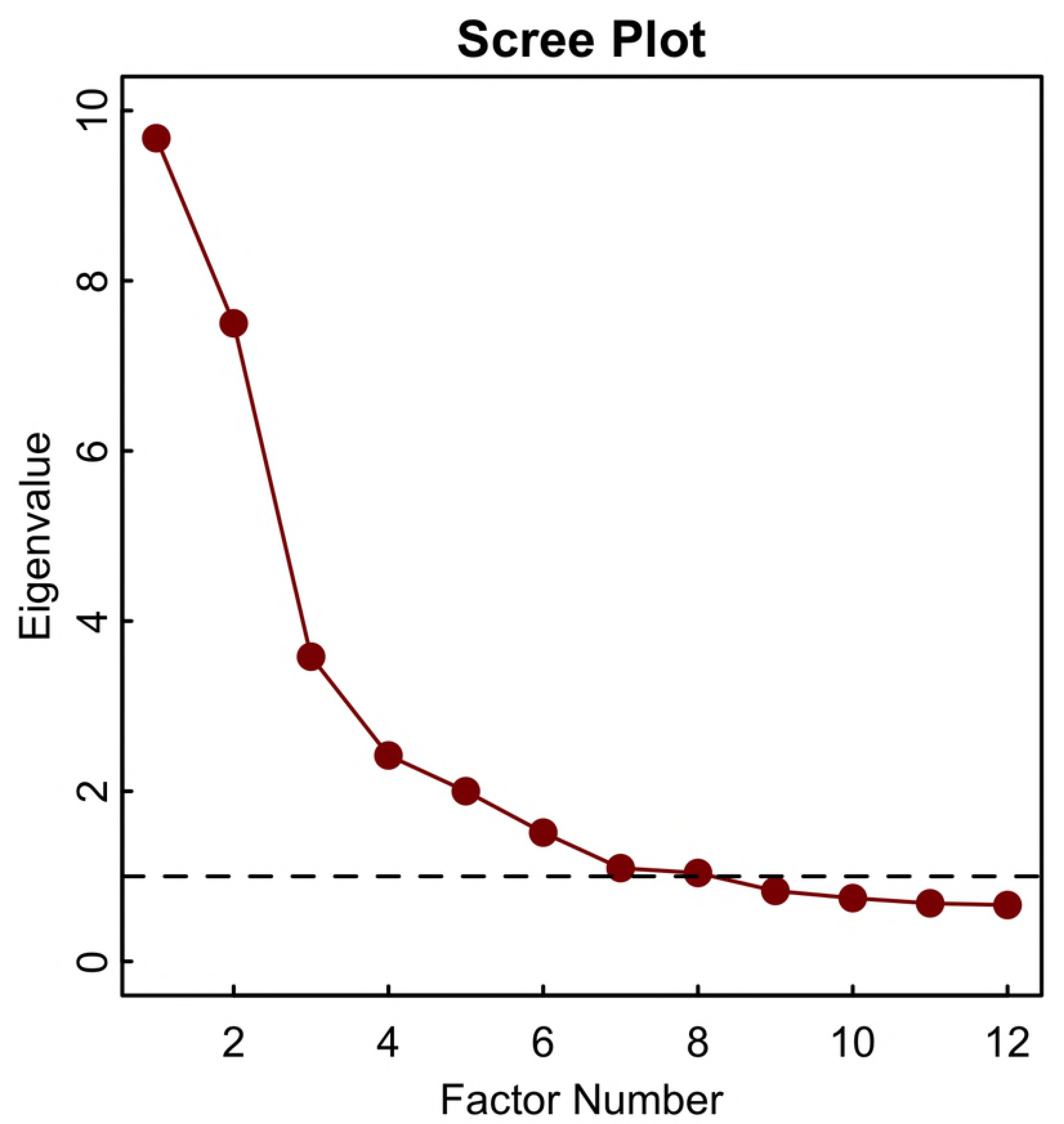

**Fig S6A.**
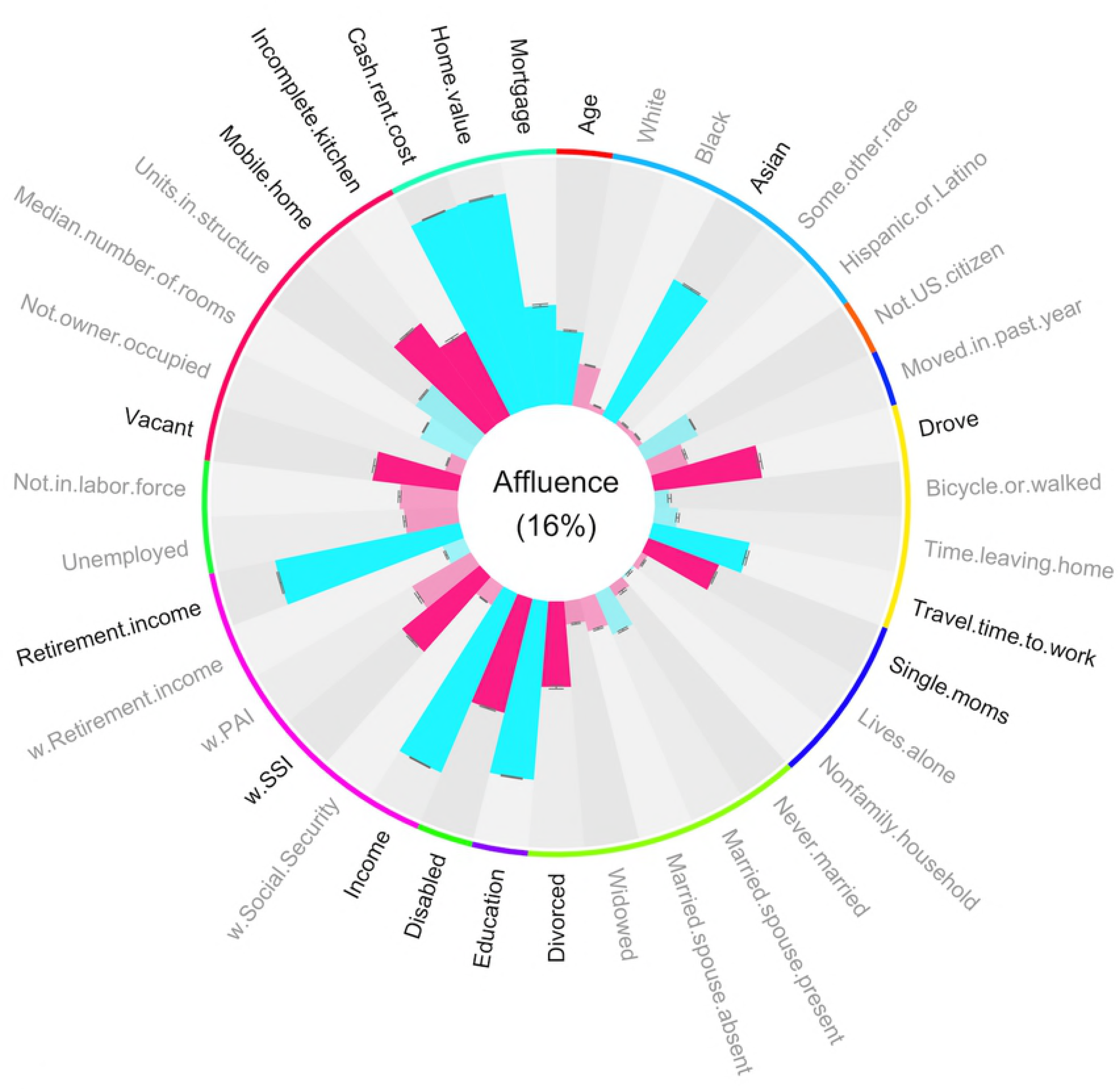

**Fig S6B.**
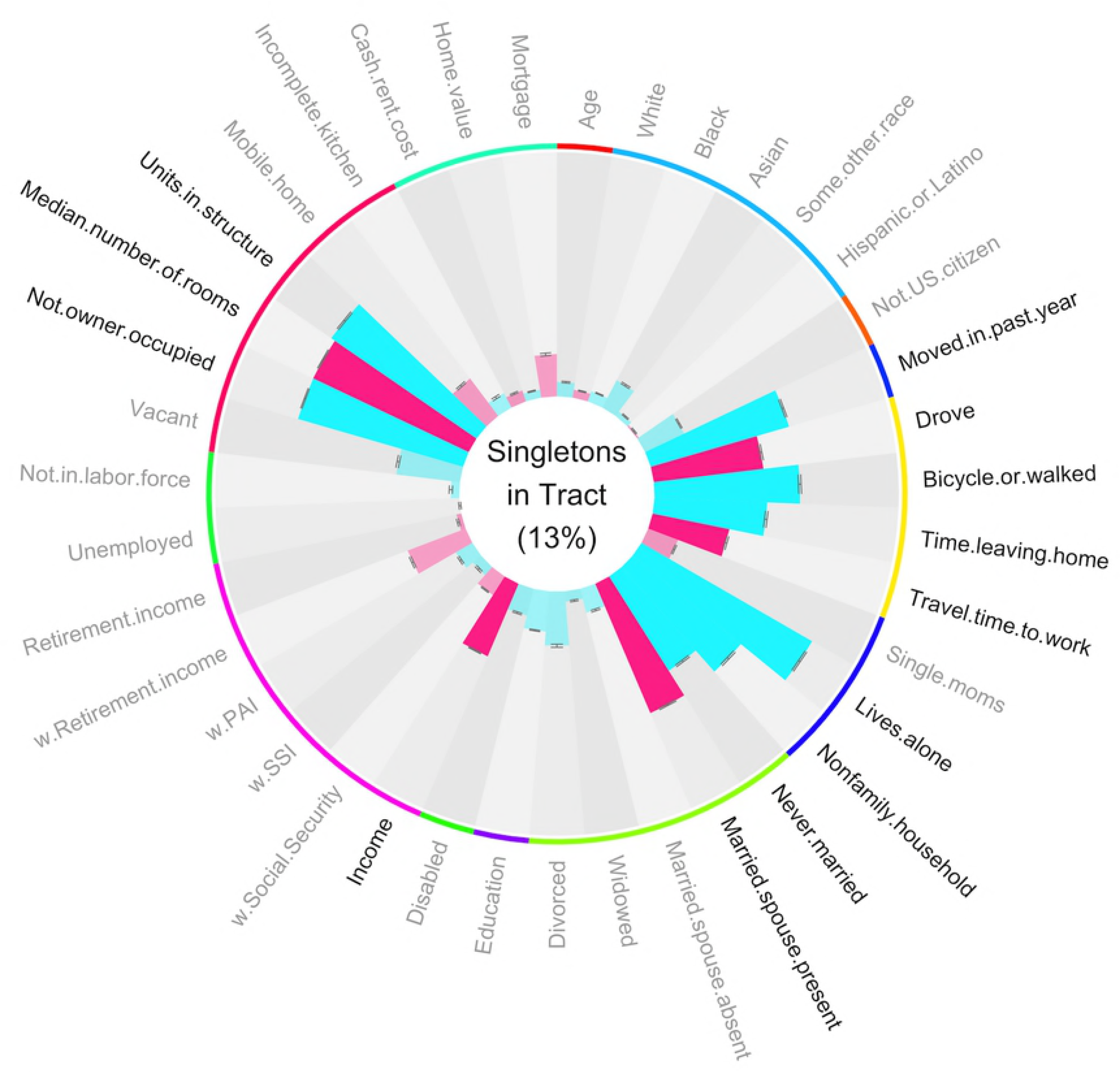

**Fig S6C.**
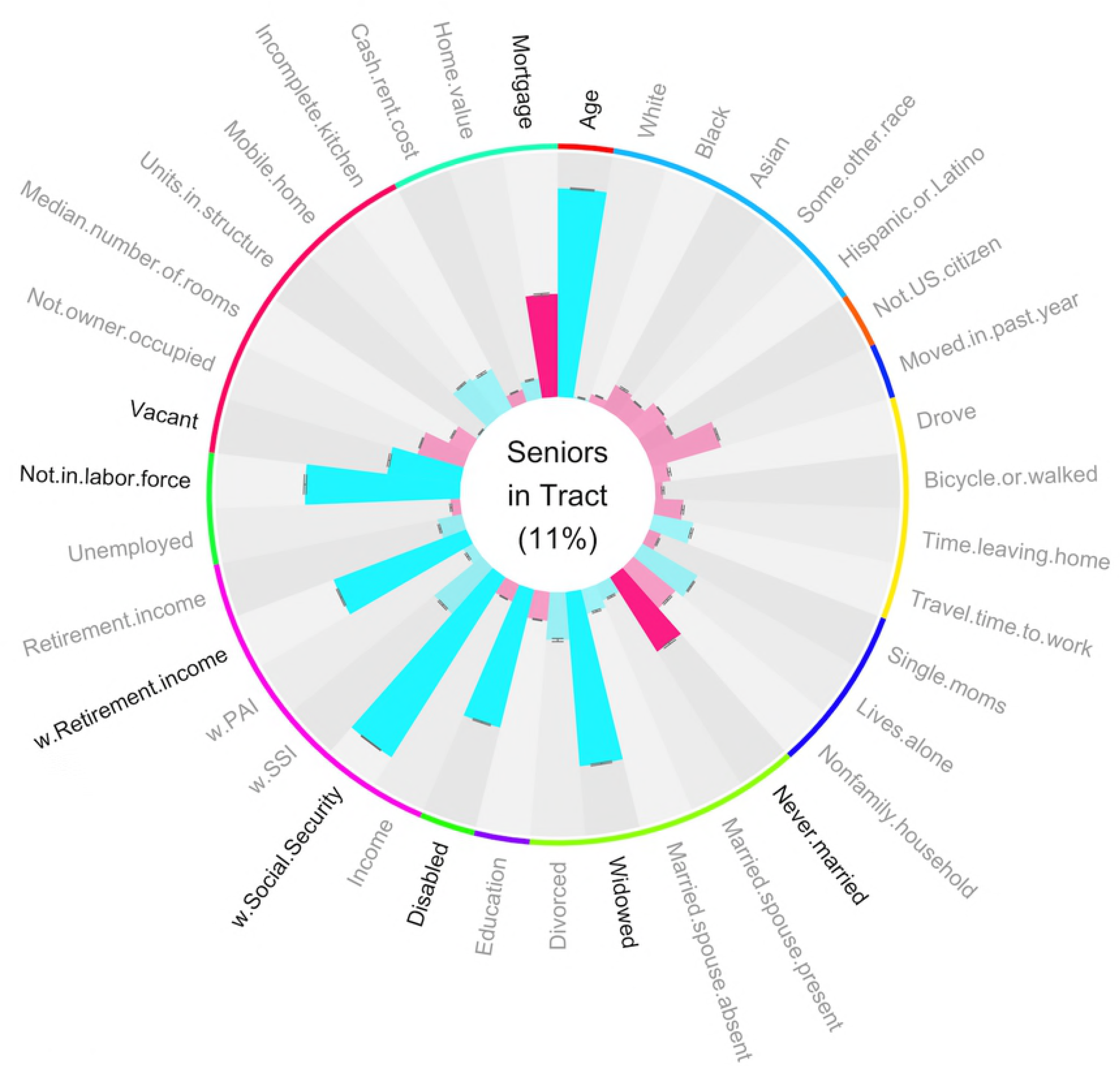

**Fig S6D.**
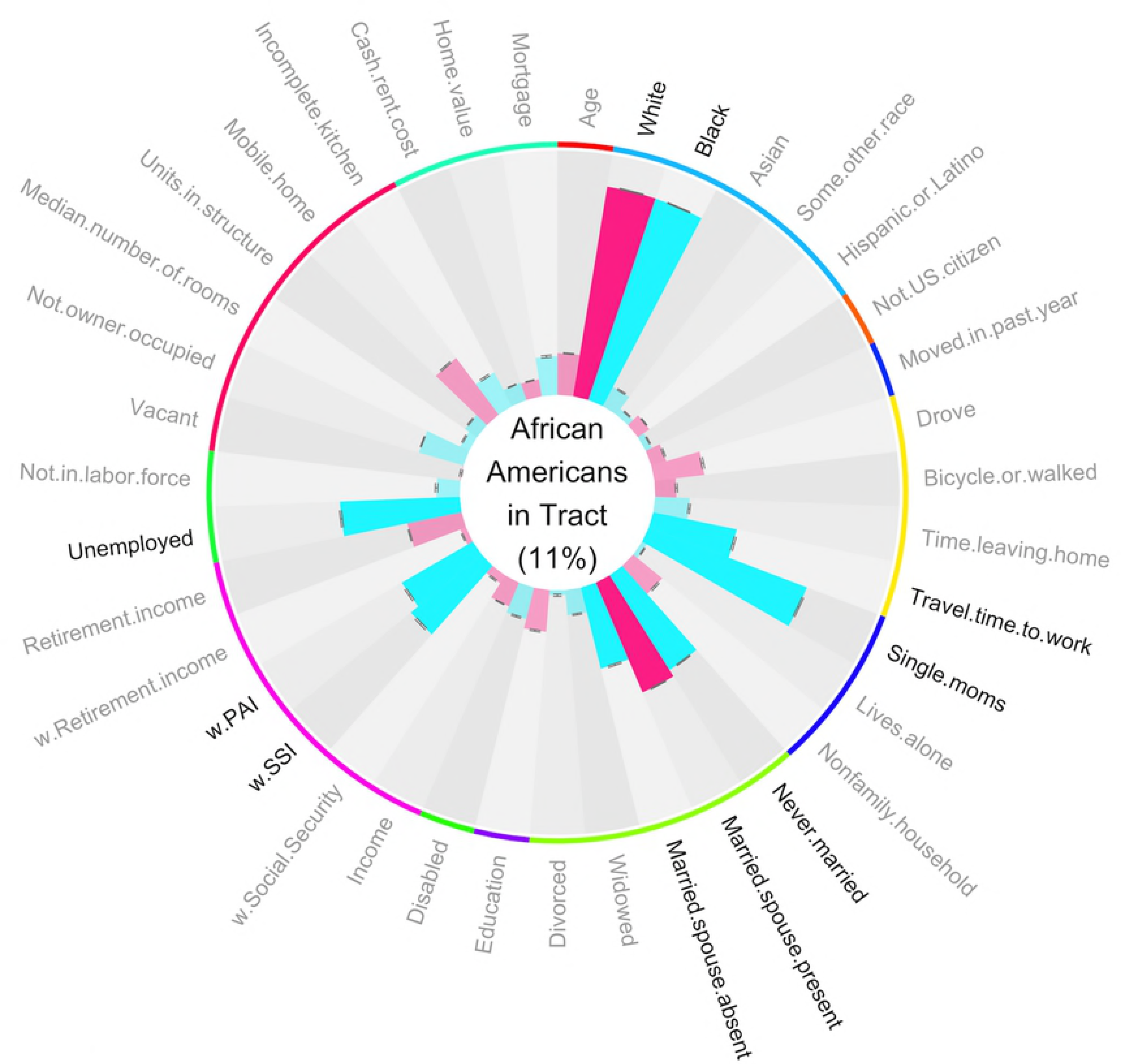

**Fig S6E.**
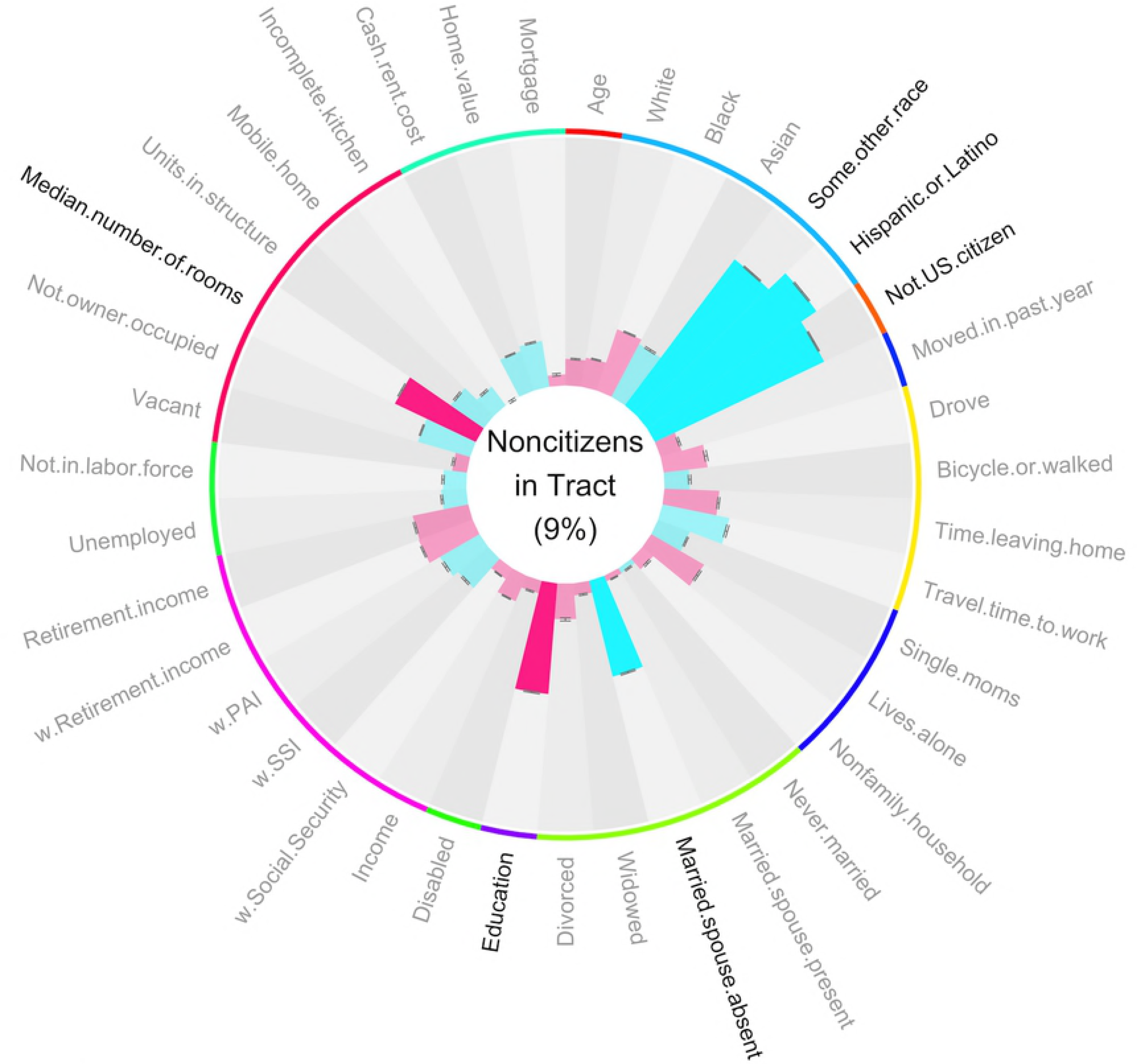

**Fig S7.**
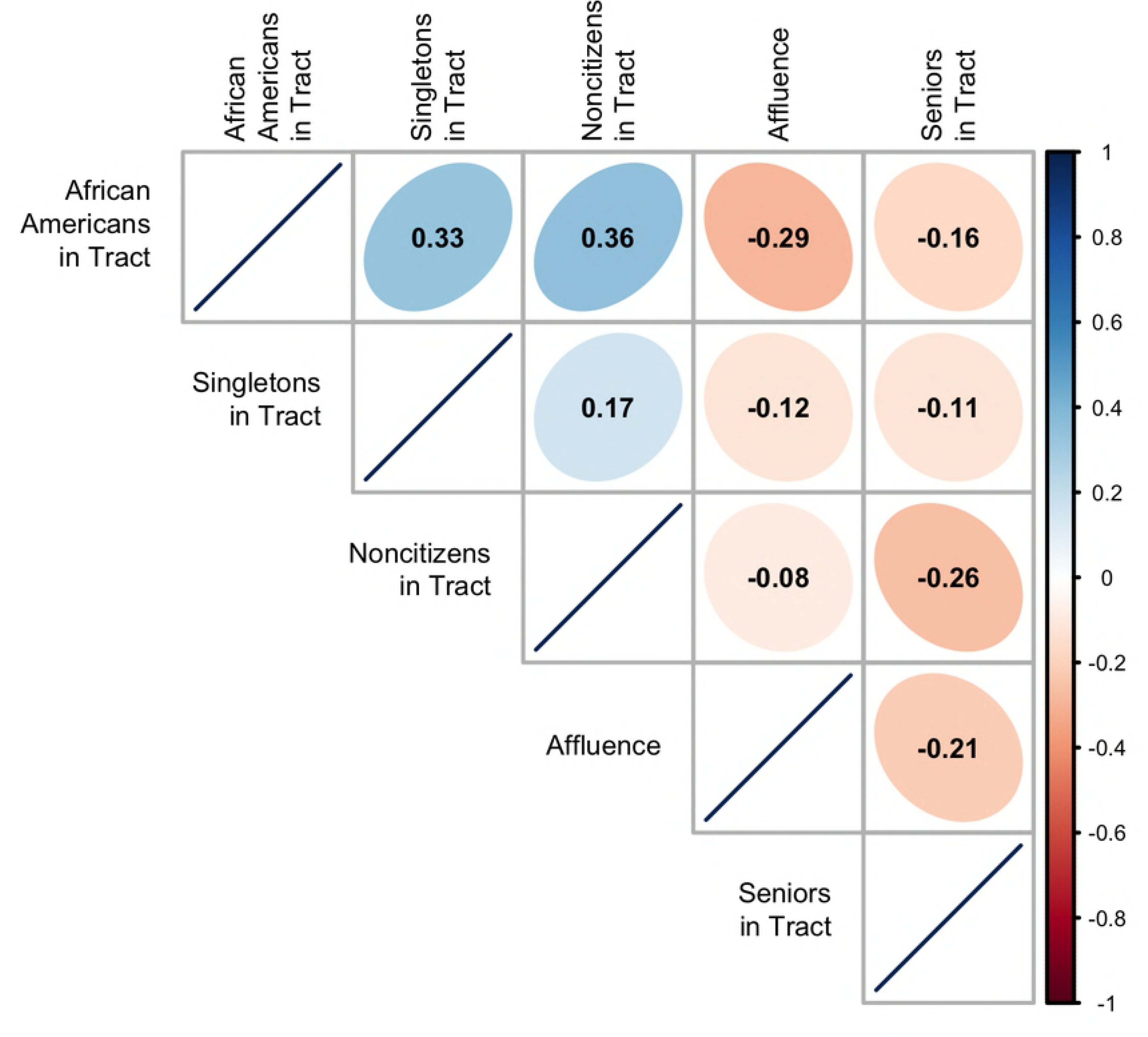

**Fig S8.**
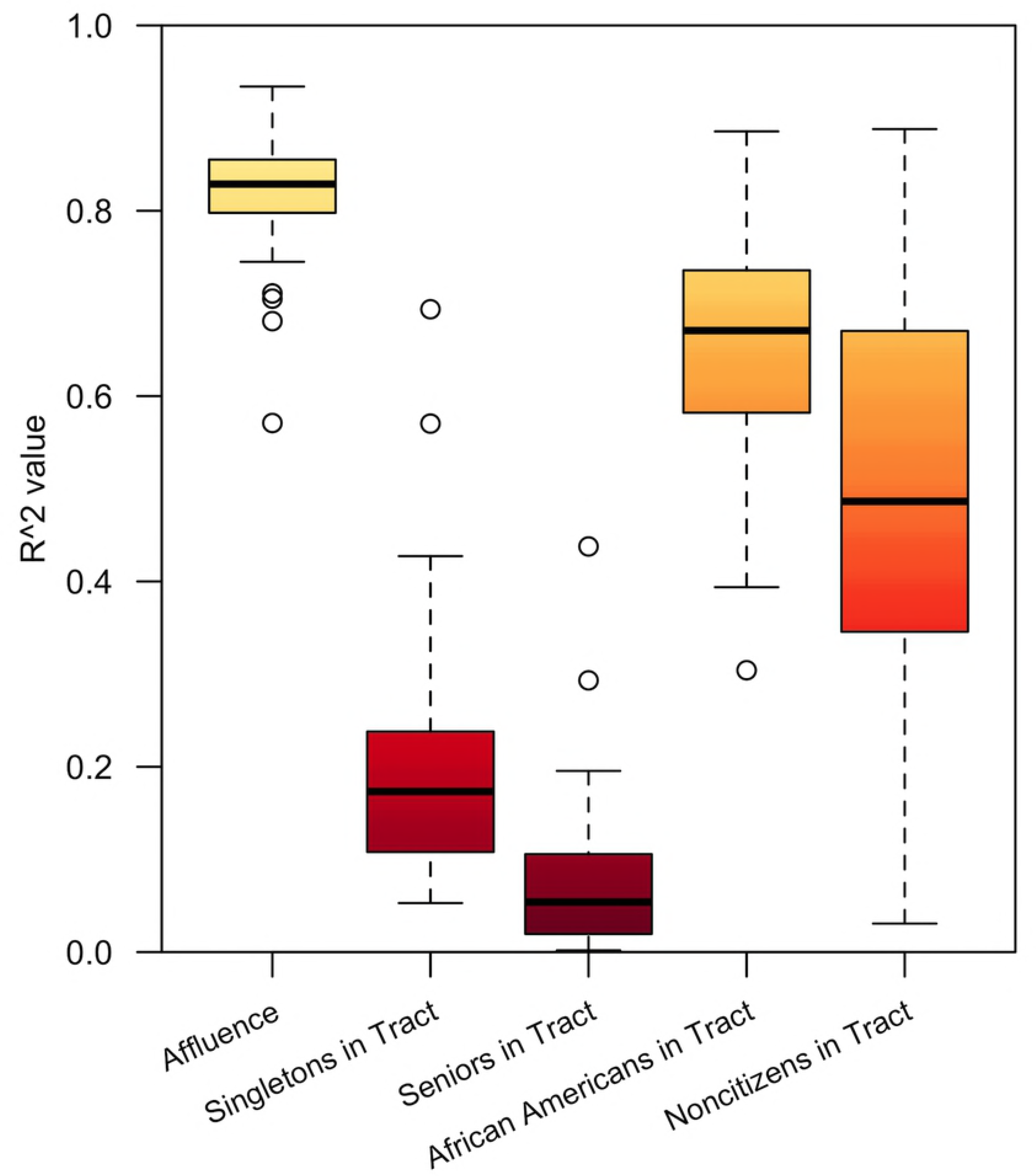

**Fig S9.**
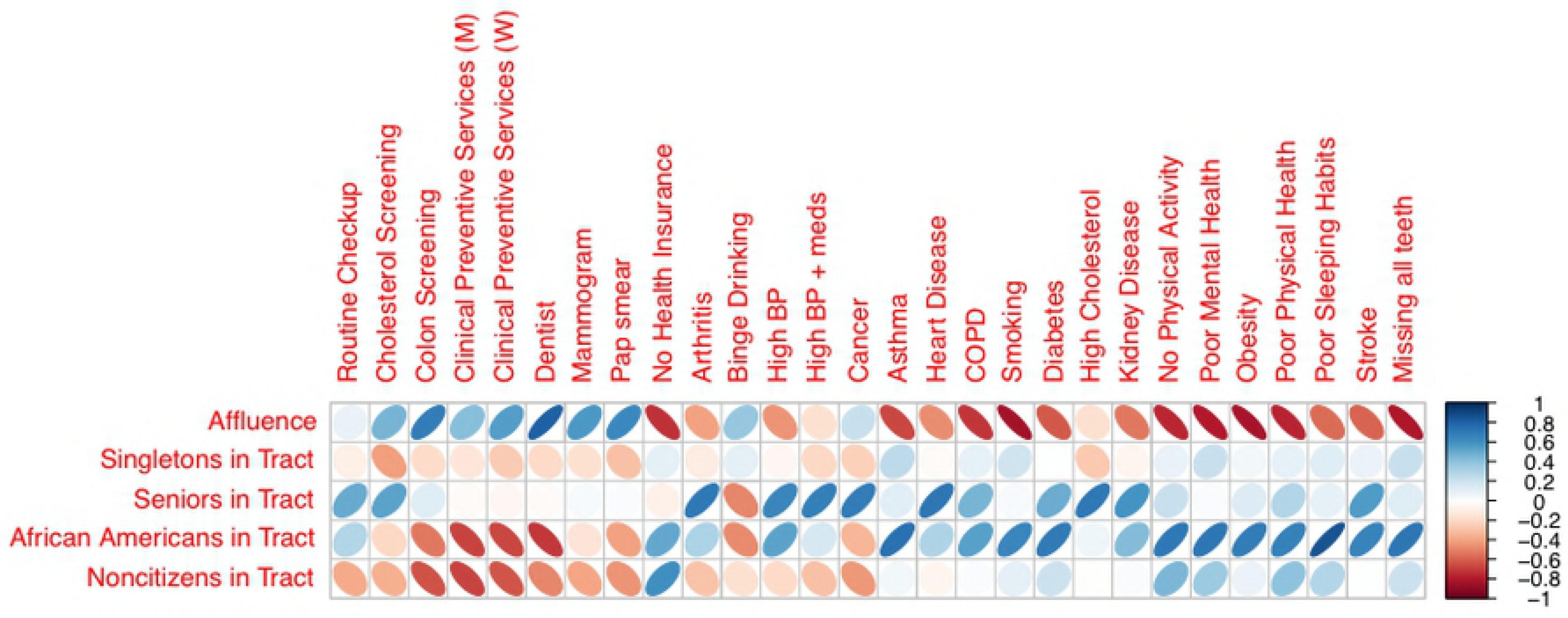

## Disclosures and acknowledgments

None of the authors have conflicting equity ownership, profit-sharing agreements, royalties, patents. Mrs. Forthman and Drs. Yeh, Kuplicki, and Paulus report no competing interests.

